## Supplementary Tables and Figures for "Vaccination against H5 HP influenza virus leads to persistent immune response in wild king penguins"

**Supplementary material**

**Supplementary Table 1: Cox proportional hazards model comparing chick survival between control (reference) and vaccinated treatment groups.** Hazard ratio (HR) = 1.50 (95% confidence interval [CI]: 0.64–3.53), *p* = 0.35, indicating no significant difference in risk between groups. SE = standard error; z = Wald test statistic.

| Variable | Coefficient (log HR) | HR | 95% CI (HR) | SE | z | *p*-value |
|---|---|---|---|---|---|---|
| Treatment [Control] | 0.43 | 1.50 | [0.64-3.53] | 0.44 | 0.93 | 0.353 |

**Supplementary Table 2: Anti-H5 and anti-NP antibody levels in king penguin chicks measured in plasma samples collected during the experimental saRNA H5 vaccination trial.** Antibody levels were obtained using ELISA assays, and were expressed either as inhibition values (competitive ELISA, H5-cELISA and NP-cELISA), or S/P values (indirect ELISA, H5-iELISA). Mean values and associated standard errors (SE) were reported for each treatment group (vaccinated and control), and for each sampling point (Day). Sampling sizes (n) refer to the number of plasma samples analysed with the corresponding ELISA assay.

| Day | | Anti-H5 antibody levels | | Anti-NP antibody levels |
|---|---|---|---|---|
| | | H5-cELISA Mean (inhibition value) ± SE | H5-iELSA Mean (S/P ratio) ± SE | NP-cELISA Mean (inhibition value) ± SE |
| 0 Primo-injection | Vaccinated | 0.96 ± 0.012 n=30 | 0.15 ± 0.046 n=30 | 0.85 ± 0.017 n=30 |
| | Control | 0.99 ± 0.0078 n=20 | 0.11 ± 0.0052 n=20 | 0.84 ± 0.021 n=20 |
| 37 Booster | Vaccinated | 0.72 ± 0.035 n=30 | 0.31 ± 0.050 n=30 | 0.83 ± 0.019 n=30 |
| | Control | 0.94 ± 0.012 n=19 | 0.15 ± 0.0078 n=19 | 0.81 ± 0.015 n=19 |
| 51 | Vaccinated | 0.055 ± 0.0073 n=30 | 4.34 ± 0.18 n=30 | 0.85 ± 0.019 n=30 |
| | Control | 0.95 ± 0.012 n=19 | 0.15 ± 0.0077 n=19 | 0.83 ± 0.018 n=19 |
| 66 | Vaccinated | 0.069 ± 0.012 n=30 | 4.67 ± 0.20 n=29 | 0.85 ± 0.022 n=29 |
| | Control | 0.97 ± 0.011 n=19 | 0.18 ± 0.010 n=19 | 0.87 ± 0.016 n=19 |
| 86 | Vaccinated | 0.088 ± 0.018 n=30 | 4.55 ± 0.23 n=30 | 0.85 ± 0.021 n=29 |
| | Control | 0.95 ± 0.011 n=19 | 0.25 ± 0.046 n=19 | 0.85 ± 0.022 n=19 |
| 164 | Vaccinated | 0.17 ± 0.036 n=29 | 3.72 ± 0.28 n=29 | 0.86 ± 0.027 n=29 |
| | Control | 0.93 ± 0.013 n=19 | 0.27 ± 0.042 n=19 | 0.86 ± 0.027 n=19 |
| 252 | Vaccinated | 0.22 ± 0.054 n=19 | 3.49 ± 0.39 n=19 | 0.84 ± 0.023 n=19 |
| | Control | 0.95 ± 0.013 n=10 | 0.64 ± 0.13 n=10 | 0.86 ± 0.013 n=19 |

**Supplementary Table 3: Pairwise contrasts between control and vaccinated groups at each sampling day, estimated from a linear mixed-effects model based on H5 competitive ELISA results.** Estimated marginal means (Estimate) refer to the difference in anti-H5 antibody titres measured using H5 competitive ELISA assay (inhibition values, H5-cELISA), between treatment groups, with associated Standard Error (SE), degrees of freedom (df), and *t*-ratio (Estimate/SE). Reported *p*-values are Tukey-adjusted for multiple comparisons. Negative values indicate lower values in the control group compared to vaccinated individuals. Significant effects are highlighted in bold: a significant difference between treatment groups is detected from day 37 onward ($p < 0.0001$).

| Day | H5 competitive ELISA assay | | | | |
|---|---|---|---|---|---|
| | Estimate | SE | df | *t*-ratio | *p*-value |
| 0<br>Primo-injection | -0.0394 | 0.0327 | 204 | -1.205 | 0.2295 |
| 37<br>Booster | -0.2169 | 0.0331 | 208 | -6.548 | **<0.0001** |
| 51 | -0.8974 | 0.0331 | 207 | -27.076 | **<0.0001** |
| 66 | -0.9008 | 0.0331 | 207 | -27.177 | **<0.0001** |
| 86 | -0.8654 | 0.0331 | 207 | -26.111 | **<0.0001** |
| 164 | -0.7571 | 0.0333 | 209 | -22.723 | **<0.0001** |
| 252 | -0.7329 | 0.0420 | 281 | -17.448 | **<0.0001** |

**Supplementary Table 4: Pairwise contrasts of estimated marginal means based on corrected seroprevalence between days, within the control group, resulting from H5 competitive ELISA assay.** Estimates refer to differences in corrected seroprevalence of control birds between day 0 and subsequent sampling times, resulting from anti-H5 antibody titres measured with H5 competitive ELISA assay. Associated standard errors (SE), degrees of freedom (df) and *t*-ratio (Estimate/SE) are detailed for each pairwise contrast. Reported *p*-values are Tukey-adjusted for multiple comparisons. No significant differences were detected, indicating that anti-H5 antibody levels in control individuals remained stable over time.

| Contrast | H5 competitive ELISA assay | | | | |
|---|---|---|---|---|---|
| | Estimate | SE | df | *t*-ratio | *p*-value |
| Day 0 – 37 | -0.060 | 0.030 | 265 | -1.992 | 0.4225 |
| Day 0 – 51 | -0.048 | 0.030 | 265 | -1.588 | 0.6904 |
| Day 0 – 66 | -0.031 | 0.030 | 265 | -1.021 | 0.9489 |
| Day 0 – 86 | -0.047 | 0.030 | 265 | -1.553 | 0.7125 |
| Day 0 – 164 | -0.070 | 0.030 | 265 | -2.305 | 0.2454 |
| Day 0 – 252 | -0.048 | 0.037 | 270 | -1.294 | 0.8541 |

**Supplementary Table 5: Parameter estimates and associated standard errors, *t*-values and *p*-values for the reduced non-linear model describing the dynamics of antibody levels of control and vaccinated chicks, using H5 competitive ELISA results.** The model is associated with the following equation (db=dc=0):

$$f(days)=\frac{(a_{control}+da \cdot treatment) \cdot days^2}{b_{control}+c_{control} \cdot days+days^2}$$

where days is the number of days post primo-injection (exact number of days) and treatment is a binary variable indicating group identity (treatment=0 for control and treatment=1 for vaccinated individuals). Significant effects (p<0.01) are shown in bold.

| | Estimate | Standard Error | *t*-value | Pr(>\|t\|) |
|---|---|---|---|---|
| $a_{control}$ | 0.032 | 0.009 | 3.629 | **0.000331 *** ** |
| da | 0.488 | 0.023 | 21.463 | **< 2e-16 *** ** |
| $b_{control}$ | 3262.91 | 165.7 | 19.696 | **< 2e-16 *** ** |
| $c_{control}$ | -79.04 | 3.85 | -20.552 | **< 2e-16 *** ** |

**Supplementary Table 6: Pairwise contrasts between control and vaccinated groups at each sampling day, estimated from a linear mixed-effects model based on H5 indirect ELISA results.** Estimated marginal means (Estimate) refer to the difference in anti-H5 antibody titres measured using H5 direct ELISA assay (S/P values, H5-iELISA), between treatment groups, with associated Standard Error (SE), degrees of freedom (df), and *t*-ratio (Estimate/SE). Reported *p*-values are Tukey-adjusted for multiple comparisons. Negative values indicate lower values in the control group compared to vaccinated individuals. Significant effects are highlighted in bold: a significant difference between treatment groups is detected from day 51 onward ($p < 0.05$).

| Day | H5 indirect ELISA assay | | | | |
|---|---|---|---|---|---|
| | Estimate | SE | df | *t*-ratio | *p*-value |
| 0<br>Primo-injection | -0.0464 | 0.247 | 163 | -0.188 | 0.851 |
| 37<br>Booster | -0.2062 | 0.249 | 165 | -0.828 | 0.409 |
| 51 | -4.1870 | 0.250 | 167 | -16.752 | **<0.0001** |
| 66 | -4.5135 | 0.251 | 168 | -17.974 | **<0.0001** |
| 86 | -4.2985 | 0.250 | 167 | -17.198 | **<0.0001** |
| 164 | -3.4506 | 0.251 | 168 | -13.743 | **<0.0001** |
| 252 | -2.9765 | 0.309 | 250 | -9.627 | **<0.0001** |

**Supplementary Table 7: Pairwise contrasts of estimated marginal means based on corrected seroprevalence between, within the control group, resulting from H5 indirect ELISA assay.** Estimates refer to differences in corrected seroprevalence of control birds between day 0 and subsequent sampling times, resulting from anti-H5 antibody titres measured with H5 indirect ELISA assay. Associated standard errors (SE), degrees of freedom (df) and *t*-ratio (Estimate/SE) are detailed for each pairwise contrast. Reported *p*-values are Tukey-adjusted for multiple comparisons. No significant differences were detected, indicating that anti-H5 antibody levels in control individuals remained stable over time.

| Contrast | H5 indirect ELISA assay | | | | |
|---|---|---|---|---|---|
| | Estimate | SE | df | *t*-ratio | *p*-value |
| Day 0 – 37 | -0.048 | 0.213 | 265 | -0.225 | 1.000 |
| Day 0 – 51 | -0.049 | 0.213 | 265 | -0.231 | 1.000 |
| Day 0 – 66 | -0.074 | 0.213 | 265 | -0.346 | 0.9999 |
| Day 0 – 86 | -0.141 | 0.213 | 265 | -0.663 | 0.994 |
| Day 0 – 164 | -0.166 | 0.213 | 265 | -0.779 | 0.987 |
| Day 0 – 252 | -0.516 | 0.262 | 269 | -1.968 | 0.437 |

**Supplementary Table 8: Parameter estimates and associated standard errors, *t*-values, and *p*-values for the reduced non-linear model describing antibody levels over time of control and vaccinated chicks, using H5 indirect ELISA results.** The model is associated with the following equation (db=dc=0):

$$f(days)=\frac{(a_{control}+da \cdot treatment) \cdot days^2}{b_{control}+c_{control} \cdot days+days^2}$$

where days is the number of days post primo-injection and treatment is a binary variable indicating group identity (treatment=0 for control and treatment=1 for vaccinated individuals). Significant effects ($p$<0.05) are shown in bold.

| | Estimate | Standard Error | *t*-value | Pr(>\|t\|) |
|---|---|---|---|---|
| $a_{control}$ | 0.12201 | 0.052 | 2.326 | **0.0207 *** ** |
| da | 1.87257 | 0.134 | 14.026 | **< 2e-16 *** ** |
| $b_{control}$ | 4131.56 | 235.245 | 17.563 | **< 2e-16 *** ** |
| $c_{control}$ | -99.89 | 4.82 | -20.726 | **< 2e-16 *** ** |

**Supplementary Table 9: Plasma seroneutralisation activity of saRNA H5 vaccinated king penguin chicks challenged by a HP H5N1 virus.** Mean seroneutralisation titre (Mean SNT titre) and associated standard errors (SE), estimated marginal means (EM Mean ± SE) and 95% confidence intervals (95% CI) were calculated for each treatment group (vaccinated and control), on chicks sampled after the boost-injection (50 days post primo-injection), and on the same birds recaptured before fledging (252 days post primo-injection). Values were obtained from a linear mixed-effects model (log(titre) ~ treatment * days + (1|bird ID)). Sampling sizes (n) refer to the number of plasma samples analysed.

| Days | Plasma HP H5N1 seroneutralisation activity | | | | |
|---|---|---|---|---|---|
| | Mean SNT titre ± SE | | EM Mean ± SE (log transformed) | 95% CI (lower-upper) | Sample size (n) |
| 51 | Control | 5.6 ± 0.98 | 2.40 ± 0.46 | 1.45 – 3.35 | n = 5 |
| | Vaccinated | 2307 ± 149 | 11.13 ± 0.32 | 10.46 – 11.80 | n = 10 |
| 252 | Control | 7.2 ± 2.3 | 2.60 ± 0.46 | 1.65 – 3.55 | n = 5 |
| | Vaccinated | 1082 ± 299 | 9.43 ± 0.32 | 8.76 – 10.10 | n = 10 |

**Supplementary Table 10: Pairwise comparison of plasma HP H5N1 seroneutralisation titre (log-transformed values) between groups (control vs vaccinated), and sampling days (50 and 250 days post primo-injection), based on estimated marginal means from the linear mixed-effects model.** Values represent estimated differences (Estimate) and associated standard errors (SE), with degrees of freedom (df), *t*-ratio and Tukey-adjusted *p*-values. Significant effects are reported in bold.

| Contrast | Plasma HP H5N1 seroneutralisation activity | | | | |
|---|---|---|---|---|---|
| | Estimate | SE | df | *t*-ratio | *p*-value |
| Control (day 51) – Vaccinated (day 51) | -8.73 | 0.561 | 23.6 | -15.558 | **<0.0001** |
| Control (day 51) – Control (day 252) | -0.20 | 0.535 | 13.0 | -0.374 | 0.9814 |
| Control (day 51) – Vaccinated (day 252) | -7.03 | 0.561 | 23.6 | -12.526 | **<0.0001** |
| Vaccinated (day 51) – Control (day 252) | 8.53 | 0.561 | 23.6 | 15.202 | **<0.0001** |
| Vaccinated (day 51) – Vaccinated (day 252) | 1.70 | 0.378 | 13.0 | 4.497 | **0.0029** |
| Control (day 252) – Vaccinated (day 252) | -6.83 | 0.561 | 23.6 | -12.170 | **<0.0001** |

**Supplementary Table 11: Type III ANOVA table based on anti-NP antibody titres measured in control and vaccinated group.** Numerator degrees of freedom (NumDF), denominator degrees of freedom (DenDF), F-value and p-values are represented for each factor tested (Effect): treatment (Group), sampling day (Day), and their interaction (Group x Day). No significant effect was detected, suggesting that anti-NP antibody levels remained stable over time, for both treatment group (all $p>0.39$).

| Effect | NumDF | DenDF | F value | *p*-value |
|---|---|---|---|---|
| Group | 1 | 48.15 | 0.0097 | 0.9221 |
| Day | 6 | 261.12 | 1.0522 | 0.3919 |
| Group × Day | 6 | 261.12 | 0.3718 | 0.8965 |

**Supplementary Table 12: Mean (± SE) number of days since injection for each grouped sampling day.** For clarity of presentation and statistical analyses, sampling days were grouped into seven categories rather than using the exact day for each individual.

| Categorical days | Mean actual days (± SE) | n |
|---|---|---|
| 0 | 0.0 ± 0.0 | 50 |
| 37 | 37.4 ± 0.07 | 49 |
| 51 | 51.4 ± 0.07 | 49 |
| 66 | 65.9 ± 0.10 | 49 |
| 86 | 86.9 ± 0.10 | 49 |
| 164 | 164.0 ± 0.10 | 48 |
| 252 | 252.0 ± 0.52 | 29 |

**Supplementary Table 13: Detailed between- and within-assay coefficients of variation (CV) for each ELISA assay**. Mean CV and associated standard error (SE) are based on values of repeated sub-samples (replicates) analysed on each ELISA plate run, and are expressed in percentage. The minimum and maximum CV (CVmin - CVmax) are indicated for each ELISA assay, and each variation (between- or within-assay).

| ELISA assay | ELISA method | Variation | Coefficient of Variation (Mean CV ± SE %) | CVmin - CVmax (%) | Number of sample replicates | Number of plates runs |
|---|---|---|---|---|---|---|
| anti-H5 | Competition | Between-assay | 3.39 ± 0.34 | 2.59 – 4.44 | 5 | 4 |
| | | Within-assay | 1.65 ± 0.60 | 0.45 – 3.94 | 5 | |
| | Indirect | Between-assay | 7.26 ± 3.1 | 1.59 - 15.8 | 5 | 5 |
| | | Within-assay | 5.81 ± 1.5 | 0.60 – 19.3 | 13 | |
| anti-NP | Competition | Between-assay | 9.17 ± 3.1 | 2.69 – 18.8 | 5 | 4 |
| | | Within-assay | 8.02 ± 1.7 | 0.60 – 17.8 | 10 | |

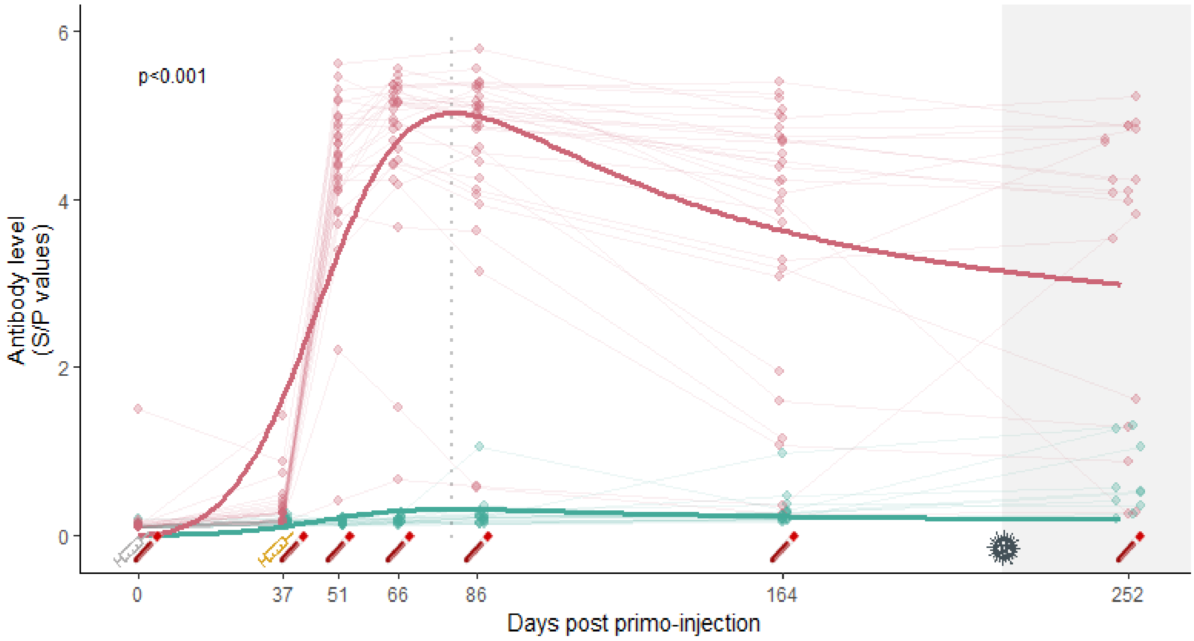


**Supplementary Figure 1: Anti-H5 *a*ntibody responses over time of king penguin chicks from saRNA H5 vaccinated (red) and control groups (blue), measured with the H5 indirect ELISA assay.** Antibody levels (expressed as [S/P ratio]) are shown for individual chicks (points), plotted at their exact sampling day from primo-injection (day 0) until fledging (day 252). Thin lines represent individual trajectories, while thicker lines indicate group-level predictions from a non-linear least-squares regression model (nlsLM) with group-specific parameters. Syringe symbols below the x-axis indicate vaccination events (grey: primo-injection at day 0; yellow: booster at day 37). Blood tube icons mark chick recaptures for blood sampling. The vertical dotted line represents the predicted timing of maximum antibody response. The horizontal dashed line shows the seropositivity threshold. The shaded grey area indicates the period after the detection of the local emergence of HPAI in October 2024 on Possession Island. The saRNA H5 vaccination induced a significant antibody response in king penguin chicks ($p$<0.001).

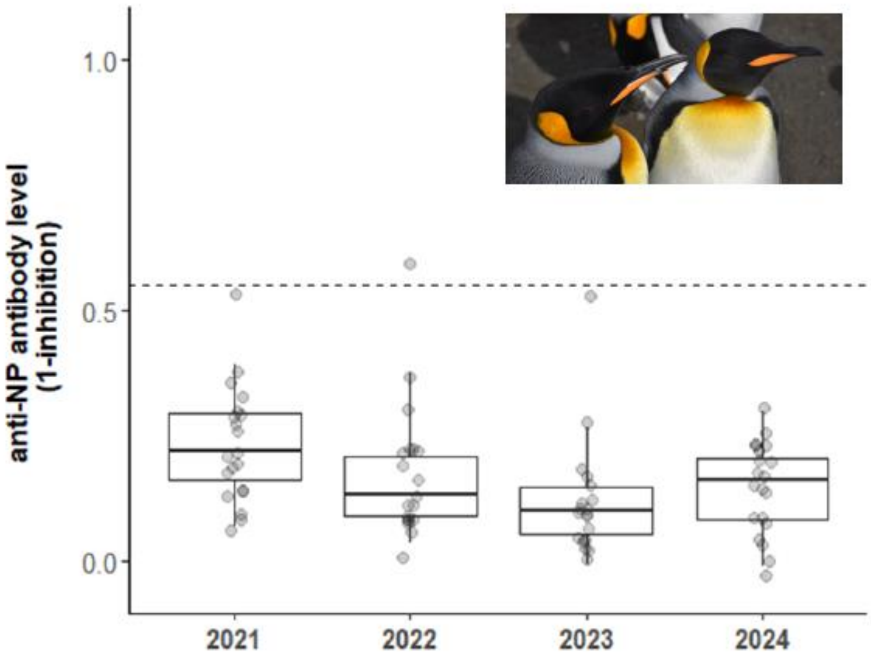


**Supplementary Figure 2: Anti-NP antibody levels of king penguin adults sampled within the colony of "Baie du Marin" (Possession Island, Crozet archipelago), from 2021 to 2024.** Antibody levels (expressed as [1-inhibition values]) of king penguin adults sampled in November 2021, 2022, 2023 and 2024 in "Baie du Marin" are represented using black dots (20 breeding adults were sampled for each field-season). The horizontal dashed line refers to the seropositivity threshold. Box plots highlight low anti-NP antibody levels expressed in king penguins suggesting that birds had not been exposed to any Influenza A viruses in this colony for the last few years (insert photograph: two adults king penguin; photo credit: T. Boulinier/CNRS/IPEV).
